# Experimental Data Connector (XDC): Integrating the Capture of Experimental Data and Metadata Using Standard Formats and Digital Repositories

**DOI:** 10.1101/2022.12.14.520467

**Authors:** Sai P. Samineni, Gonzalo Vidal, Carlos Vidal-Céspedes, Guillermo Yañez-Feliú, Timothy J. Rudge, Chris J. Myers, Jeanet Mante

## Abstract

Accelerating the development of synthetic biology applications requires reproducible experimental findings. Different standards and repositories exist to exchange experimental data and metadata. However, the associated software tools often do not support a uniform data capture, encoding, and exchange of information. A connection between digital repositories is required to prevent siloing and loss of information. To this end, we developed the Experimental Data Connector (XDC). It captures experimental data and related metadata by encoding it in standard formats and storing the converted data in digital repositories. Experimental data is then uploaded to Flapjack and the metadata to SynBioHub in a consistent manner linking these repositories. This produces complete connected experimental datasets that are exchangeable. The information is captured using a single template Excel Workbook, which can be integrated into existing experimental workflow automation processes.

**TOC Graphic:** 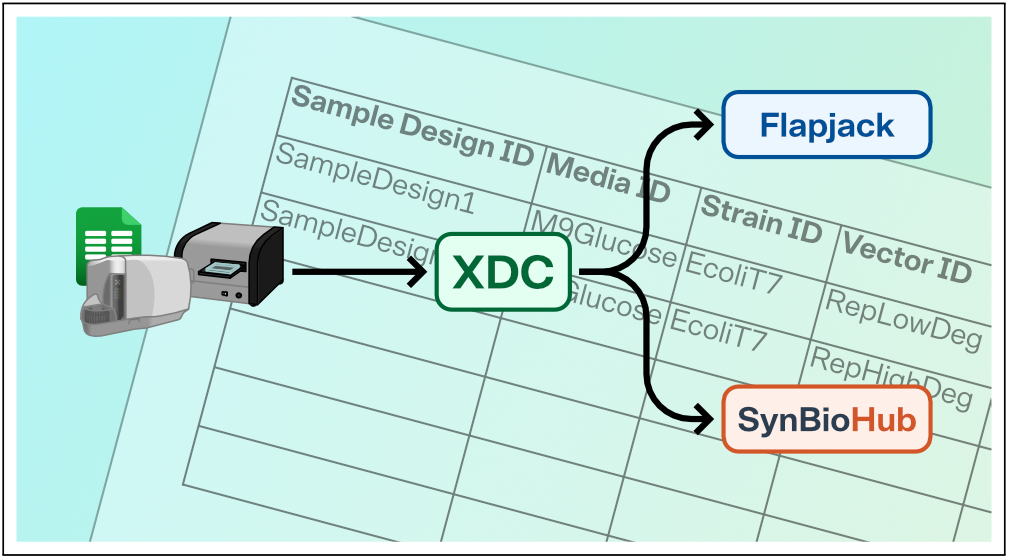

## Introduction

*Synthetic biology* (SynBio) aims to engineer biological systems in a predictable and reproducible way. To this end, software tools are needed to aid researchers in iterating the *design–build–test–learn* (DBTL) cycle. *Experimental workflow automation* (EWA) is a critical aspect of developing synthetic biology applications. In the field of SynBio, EWA promotes automation by integrating software tools to support assembly planning, protocol management, and digital data storage with well-defined sets of instructions, specific designs, materials, and machines. Through this set of software tools, EWA captures the underlying information needed to build and test genetic circuits in a reproducible manner. The use of software is essential to manage the development of complex SynBio applications which remain relevant across diverse operational contexts. Digital repositories are used to increase the reproducibility of DBTL iterations. There are several types of digital repositories, including ones focused on metadata and ones focused on experimental data. They overlap in some ways; however, they are generally not connected.

Digital repositories that support the build and test stages of EWA include SynBioHub^1^ and Flapjack.^2^ SynBioHub is a web application that provides a repository for DNA sequences, biological parts and devices, strains, experimental setup information, and other metadata (https://synbiohub.org). Flapjack is a data management system that enables researchers to store, visualize, analyze, and share genetic circuit experimental data, including measurement data and corresponding metadata (http://flapjack.rudge-lab.org/). SynBioHub and Flapjack are accessible by API frameworks to facilitate development and integration with the wider computational synthetic biology environment. Additionally, they both use the *Synthetic Biology Open Language* (SBOL).^3–5^ SBOL is a free and open-source standard for the representation and electronic exchange of information on the structural and functional aspects of biological designs.

The current workflow to upload data to SynBioHub and Flapjack is a manual, time-consuming process that requires new data imports for each new experimental context and leads to data that is not linked. This is partially due to the fact that Flapjack’s data model lacks a field to implement this connection and its frontend interface has not yet implemented a bulk upload functionality. Thus, having more than one assay implies uploading each Excel file individually to SynBioHub and Flapjack. Furthermore, plate readers from different brands produce different outputs requiring a different Flapjack parser for each one. This creating difficulties for the tool.

Here we present a new software tool, the experimental data connector (XDC), for experimental data capture using the Excel-to-SBOL Converter,^6^ Flapjack,^7^ and SynBioHub.^8^ The *XDC* uses an Excel template to capture both experimental data and metadata. Both are then converted to a uniform data representation for the SBOL standard and Flapjack’s data model. Using SBOL, users of the XDC can store experimental results with sequence, part, and other metadata information. Additionally, they can retrieve and share the data with others in a standard manner, improving reproducibility and collaboration.

## Results

The XDC, illustrated in Figure 1, provides a user-friendly workflow to capture, encode, and connect experimental data with its related metadata leveraging the established digital repositories Flapjack and SynBioHub.

**Figure 1:**
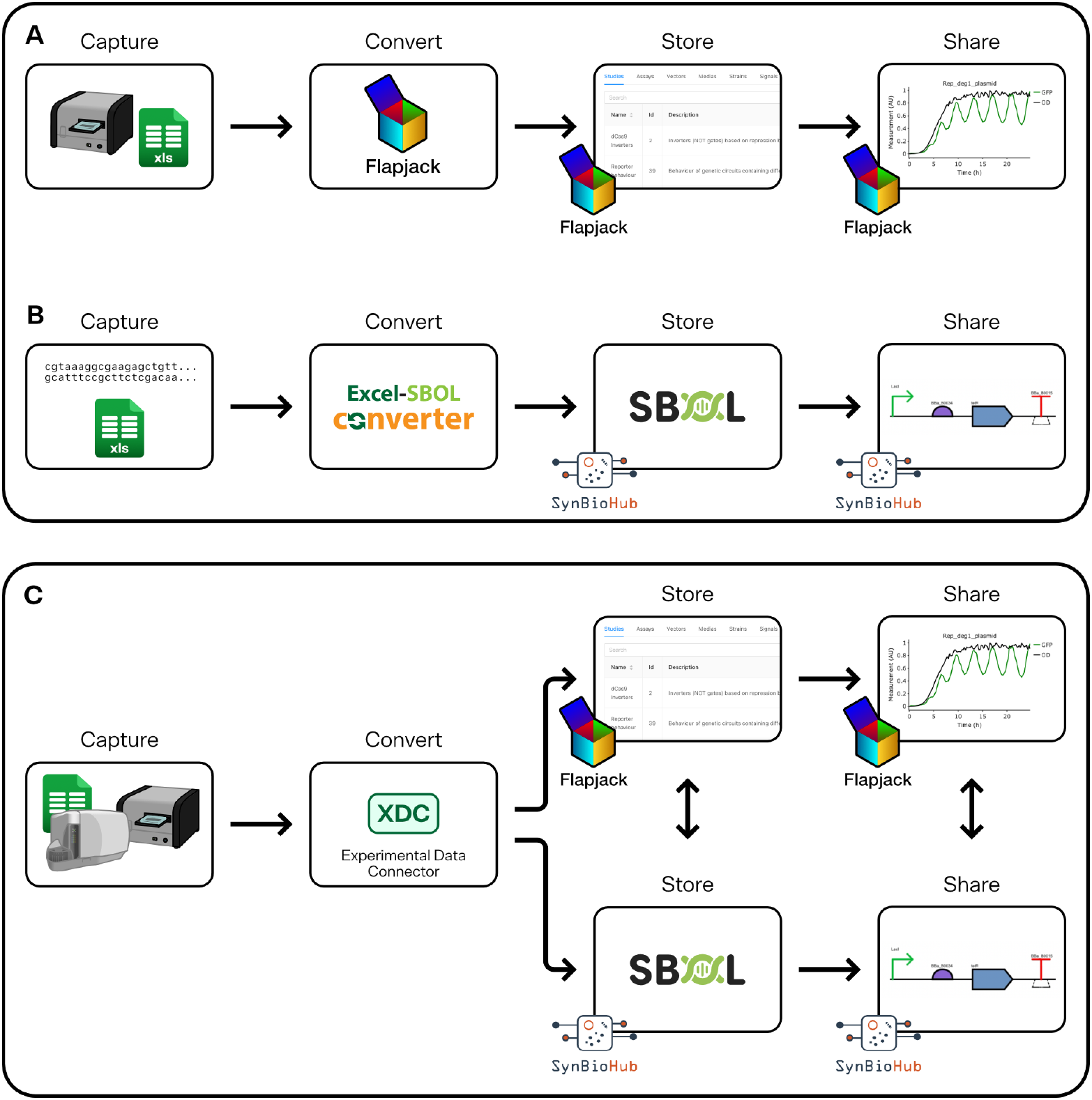
Diagram of the existing workflows (A and B) and the new workflow (C) for capturing, encoding and connecting experimental data. **A.** Workflow for storing experimental data in Flapjack. The data is captured from a plate reader, which delivers the readings in an Excel (xls) file, which is modified to incorporate the missing metadata necessary to be uploaded to Flapjack; this is then converted to the Flapjack data model and uploaded to the platform through the frontend and stored on the server. Once the data is stored, it can be viewed or shared. **B.** Workflow for storing data in SynBioHub. Genetic information is captured either through sequences obtained from text, GenBank, or by using an Excel file. This file is then converted to an SBOL file by using the Excel-to-SBOL Converter,^6^ which stores metadata in SynBioHub and subsequently is shared. **C.** Workflow using the XDC. Data can be obtained from various sources, including a plate reader and fluorescence cytometer, which are then captured by a template Excel workbook. These sheets are converted using the XDC, which generates an SBOL file with the experimental measurement data, uploads the relevant information to Flapjack with links to the SBOL objects, and uploads the SBOL representation of the metadata with links to Flapjack experimental data to SynBioHub.

The existing steps involve uploading data to Flapjack and SynBioHub as separate workflows. To store experimental data, it has to be collected in a formatted spreadsheet and uploaded to Flapjack using the frontend (Figure 1A). Then, to store the related metadata, it is necessary to capture it from a text, GenBank, or SBOL file; then, the file is later separately converted to an SBOL file as part of the SynBioHub upload process (Figure 1B). The proposed workflow streamlines this process by aggregating all the captured data into a template Excel workbook, which in turn is processed by the XDC to generate an SBOL file and upload the data to Flapjack and SynBioHub simultaneously, linking the data between the platforms in the process (Figure 1C). The XDC can be accessed via a user interface shown in Figure 2A (https://xdc.synbiohub.org/upload).

**Figure 2:**
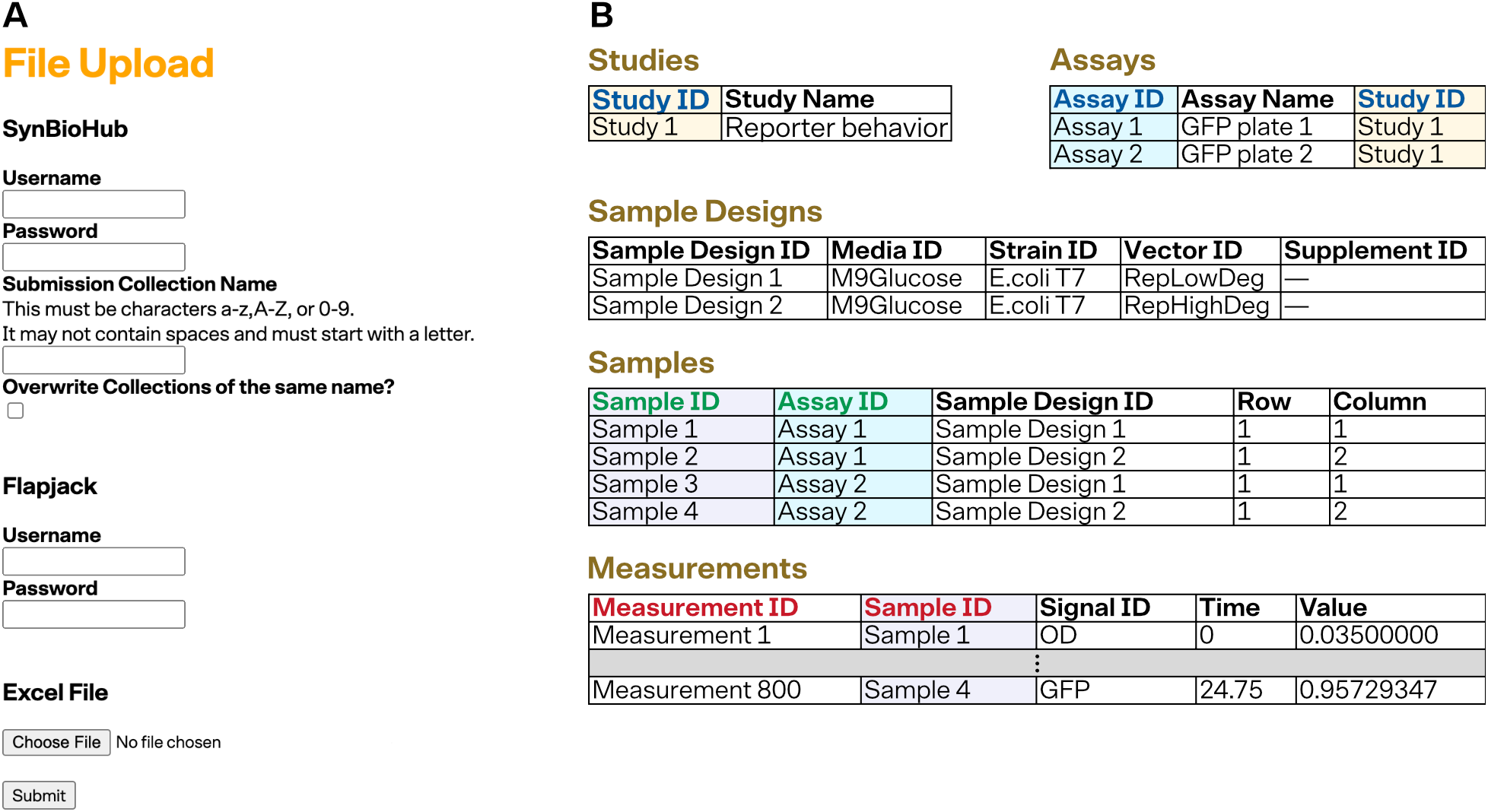
**A** The XDC front end. The XDC uses the information collected here to create the linked uploads to Flapjack and SynBioHub. **B** XDC Case Study. Showing the design of a case study experiment comprising a study with measurements every 15 minutes over a period of twenty-five hours with two repeats. For each repeat, there is one plate per repeat with one assay per plate. Each plate has two samples per assay for a total of four samples at two different sample design setups. Measuring each sample at each of the 100 time points results in 800 data points. Given this experimental setup, five different sheets are used to describe and connect information for Studies, Assays, Sample Designs, Samples, and Measurements. Note the way the sheets are linked via IDs. For example, assays identify the study they are part of using the “Study ID” column and similarly, samples to assay with “Assay ID”, and measurements to sample with “Sample ID”.

The template Excel workbook is designed to collect a wide range of experimental SynBio data (Figure 2B). It was developed as an extension of the Excel-SBOL master template.^6^ The workbook is agnostic to the equipment that generated the data allowing the use of different plate readers, flow cytometers, or any other equipment from different brands. The workbook encodes all the data fields to standardized formats using SBOL2^9^ and the Flapjack data model.^2^ If necessary, the Excel-SBOL mapping can be modified by changing the “column_definitions” sheet, and the Excel-Flapjack data mapping can be modified by changing the “FlapjackCols” sheet.

The XDC provides an improved workflow for capturing, encoding, and representing experimental data and metadata. An example of this workflow could be as follows:

1. A researcher designs an experiment to compare gene expression in repressilators with different degradation rates.
2. The researcher designs two repressilators, one with the transcription factors and GFP reporter without degradation tags and another with a degradation tag like AAV on all those proteins.
3. The researcher builds the necessary plasmids, transforms them into a chassis, and tests them by measuring OD and fluorescence on a plate reader.
4. The plate reader data is entered into the template workbook (either manually or in a semi-automated way), see Figure 2B.
5. Researchers can relate experimental measurement data to its metadata, including the conditions of the experimental study, how the DNA was built, the parts used to create the DNA, and the original developer of the parts, see Figure 2B. This Excel template workbook is provided to researchers in the XDC repository (https://github.com/SynBioDex/Xperimental-Data-Convertor).
6. The Excel workbook is uploaded to the XDC server, see Figure 2A. The XDC creates a complete linked dataset stored across Flapjack and SynBioHub

This workflow enables the analysis of the measurement data in Flapjack to be more reproducible with the capture of the study setup, sample designs, and other design data. In this way, the user is able to both identify the implementations of DNA parts employed in the study and access the underlying experimental data results through links that may be developed for measurement and analysis data in Flapjack.

## Discussion

The XDC provides a semi-automated workflow to assist researchers with the transition from build and test to the learn stage. This workflow captures data using a template Excel workbook then a software engine creates, encodes and and uploads experimental data and metadata in a standardized format. The XDC enables the use of digital repositories, Flapjack and SynBioHub, to reproduce experimental findings and improve data sharing. The user interface manages the data encoding and conversion to provide access to a wide range of researchers, particularly those with limited coding skills. Finally, the tool supports the use of the SBOL standard to capture, build and test experimental data in machine readable formats without prior knowledge of SBOL. The column definition on the Excel workbook supports the creation of new workbook versions that could encode the data into SBOL3^5^ and convert data to future versions of Flapjack or other repositories.

In the future, we plan to expand the linkage between SynBioHub and Flapjack repositories by making the Flapjack IDs link onto Flapjack viewing pages. Additionally, we intend to integrate the XDC more fully with SynBioHub by incorporating it in a submit plugin to improve user authorization as well as provide more interactivity with the templates.^10^ We envision a more connected and automated DBTL cycle in research laboratories and industrial applications. However, more tools and workflows are needed to connect the DBTL cycle fully. Automation tools like liquid handling robots and lab management systems will automatically capture relevant metadata. Automated workflows will coexist with spreadsheet metadata and experimental data capture to improve the reproducibility of experimental workflows.

## Methods

The XDC backend and frontend are written in Python. The pseudocode description of the algorithm is shown in Algorithm 1. The use of the “column_definitions” sheet is explained in the Excel-to-SBOL previous work.^6^ The “FlapjackCols” is shown in Figure 3A and takes column names and changes them to Flapjack data object names.

**Figure 3:**
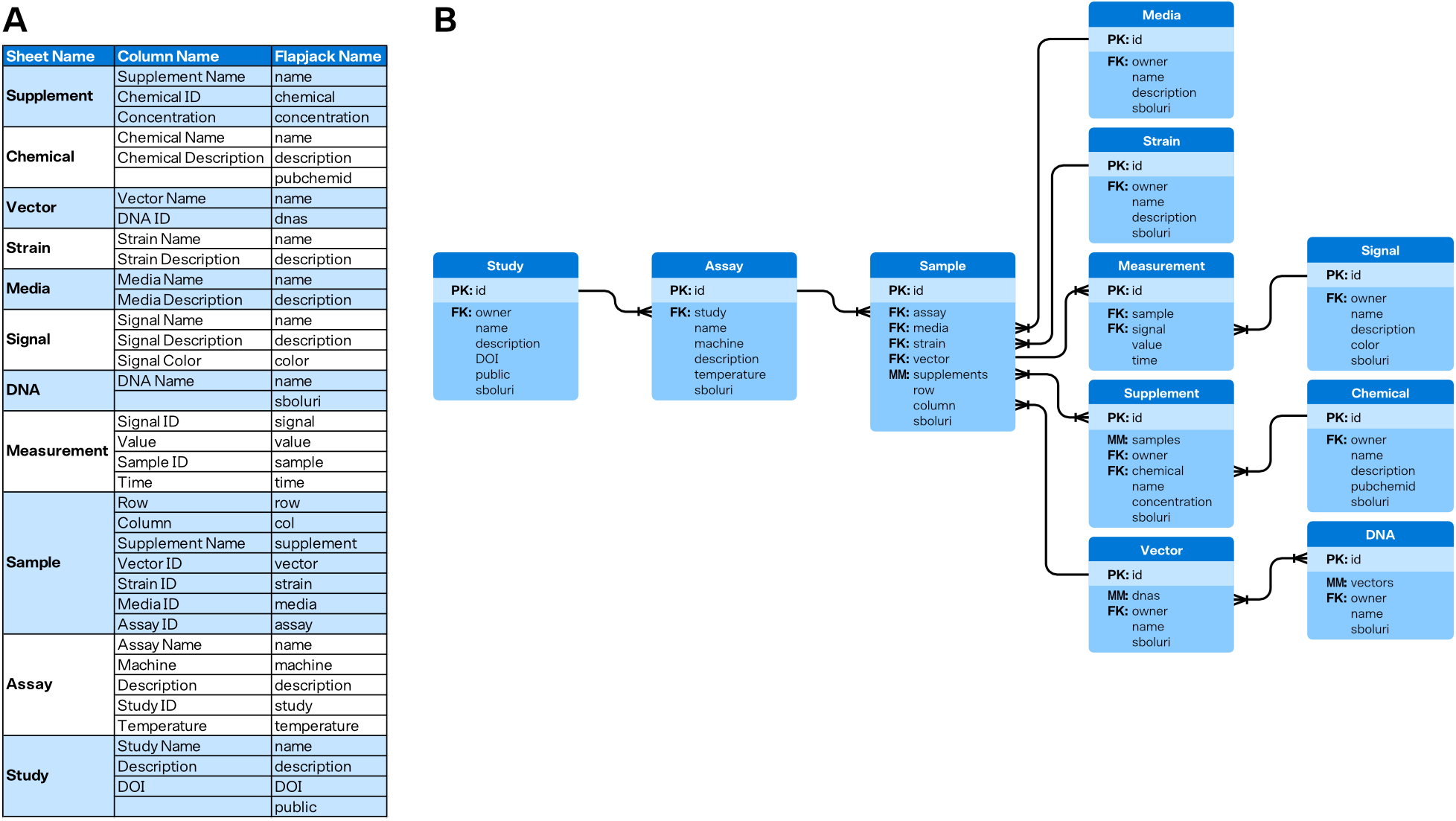
**A** FlapjackCols sheet. The Connector uses this sheet to determine how data should be uploaded to Flapjack. It provides the Sheet Name, Column Name, and the Flapjack property name it should be turned into. This allows greater flexibility in the column names while maintaining the Flapjack data model **B** The Flapjack data model. Arrows show how data types connect to each other. For example, studies contain assays.

### Algorithm 1: Experimental Data Connector

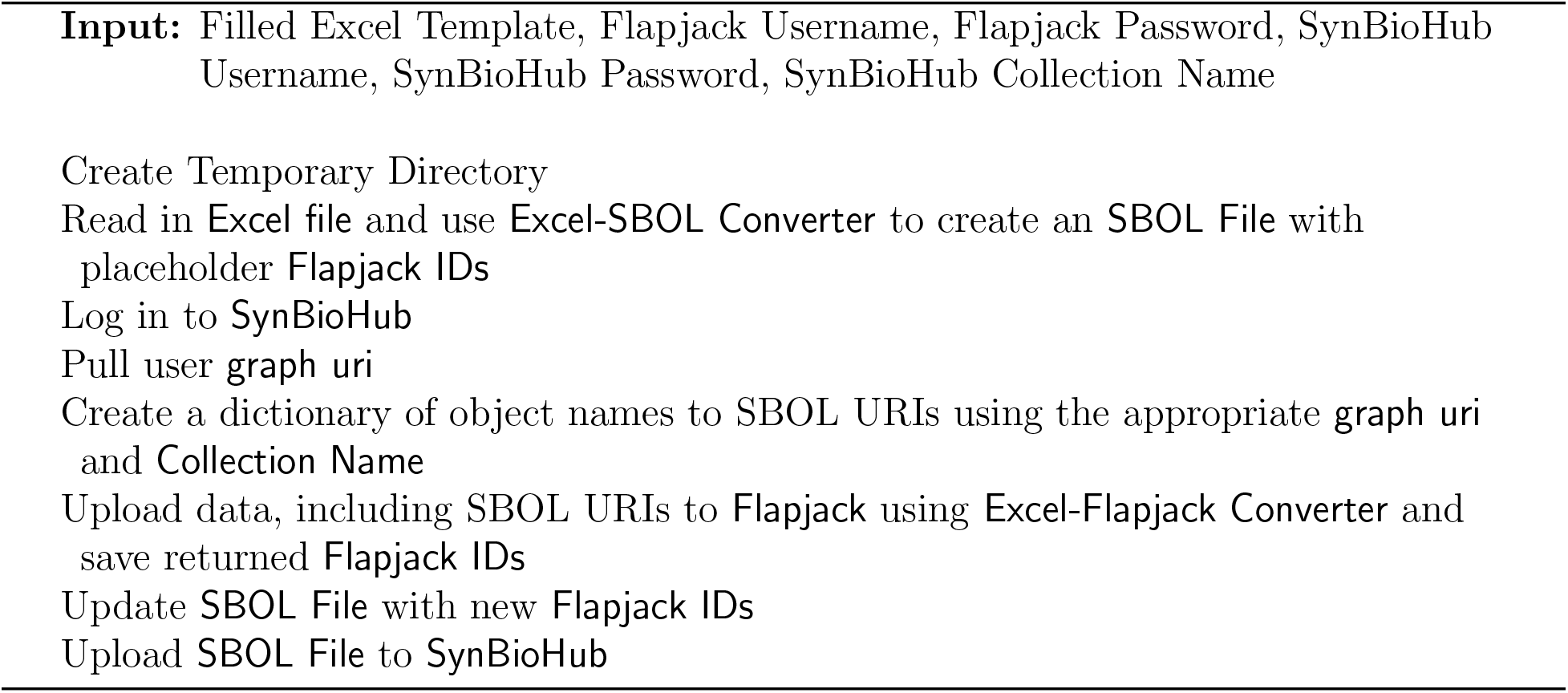

The XDC creates Flapjack objects using the excel2flapjack module (https://github.com/SynBioDex/Excel-to-Flapjack) developed in this work. The pseudocode of the excel2flapjack module is explained in Algorithm 2. The Connector creates SBOL objects using the existing Excel-to-SBOL library (https://github.com/SynBioDex/Excel-to-SBOL).

### Algorithm 2: Excel-to-Flapjack

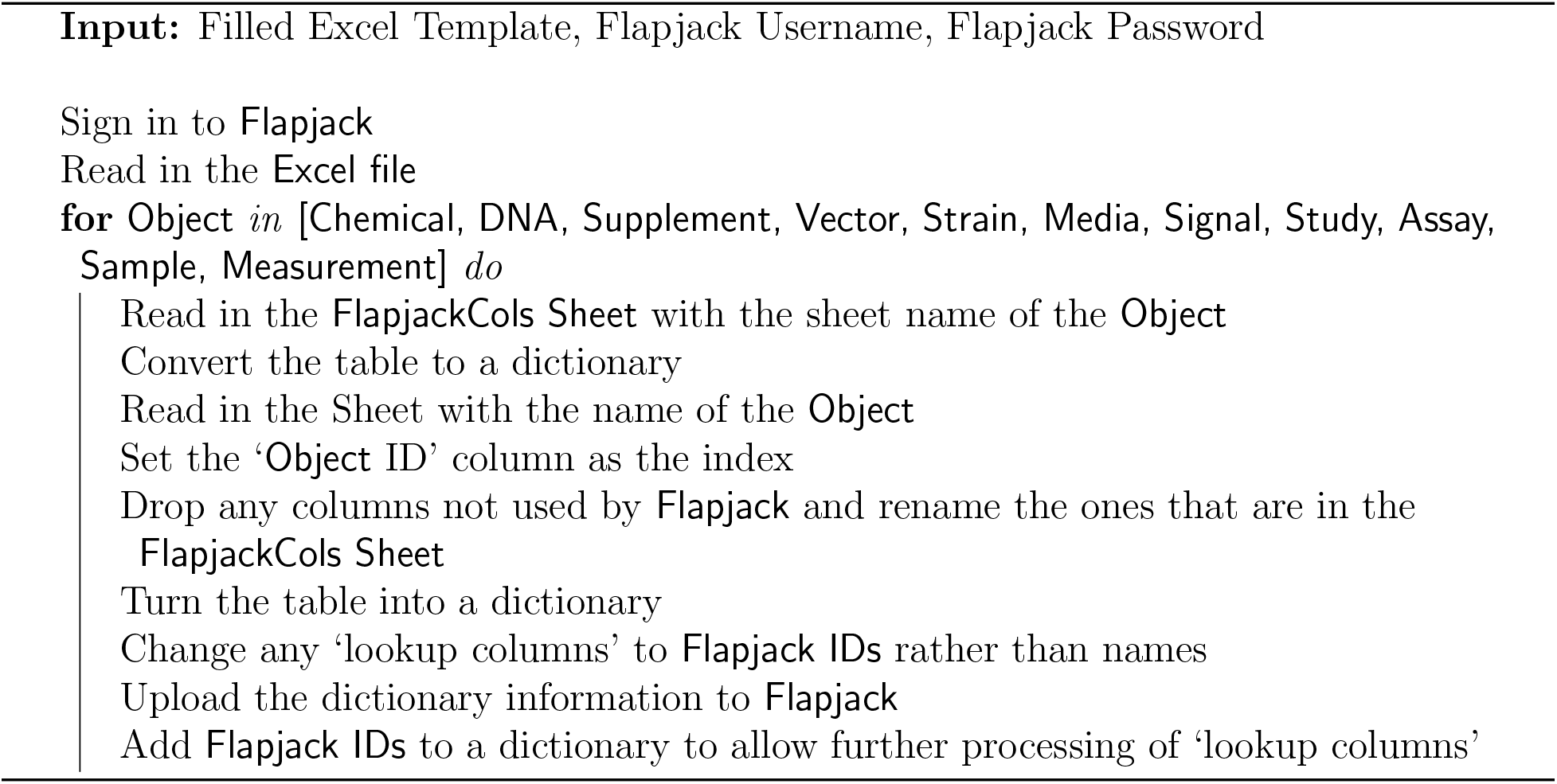

To allow the connections between SynBioHub and Flapjack, SynBioHub URI fields were added to the Flapjack data model for every object but Measurement, which is the experimental data. pyFlapjack (https://github.com/RudgeLab/pyFlapjack) was also improved in this work by extending the Create function to support overwrite prevention during the bulk upload of data. Simulated data for test and examples was generated using LOICA.^11^

## Supporting information

Example file for use with the connector

## Acknowledgement

JM, SPS, and CM are supported by the National Science Foundation under Grant Nos. 1939892 and 2231864. The Experimental Data Connector is run on a Microsoft Azure Server provided by Microsoft Research. GV, CV, GYF, and TJR are supported by the Newcastle University School of Computing. Any opinions, findings, conclusions, or recommendations expressed in this material are those of the author(s) and do not necessarily reflect the views of the funding agencies.

## Author Contributions

All authors contributed to the writing of this manuscript. SPS developed the experimental data conversion workflows supported by JM and GV. JM wrote the XDC library and front end. JM, SPS, and GV created the Excel-to-Flapjack library. GV, GYF, and CV provided support on the Flapjack API. SPS and JM created the workbook template. GV wrote the code to produce simulated data used in the Excel workbook case study. CM and TR supervised the entire project and team.

## Supporting Information Available

### Experimental Data Connector

- Server GitHub: https://github.com/SynBioDex/Xperimental-Data-Connector-Server
- Package GitHub: https://github.com/SynBioDex/Xperimental-Data-Connector
- Excel-to-Flapjack GitHub: https://github.com/SynBioDex/Excel-to-Flapjack
- Usable Instance: https://xdc.synbiohub.org/upload
- Example Template: https://github.com/SynBioDex/Xperimental-Data-Connector-Server/blob/main/xperimental_data_conncetor_v001.xlsx

### Suplemental Files

- **Excel Template for Conversion**: xperimental_data_conncetor_v001.xlsx

## References

(1) McLaughlin, J. A.; Myers, C. J.; Zundel, Z.; Misirli, G.; Zhang, M.; Ofiteru, I. D.; Goñi-Moreno, A.; Wipat, A. SynBioHub: A Standards-Enabled Design Repository for Synthetic Biology. ACS Synthetic Biology 2018, 7, 682–688.

(2) Yáñez Feliu, G.; Earle Gómez, B.; Codoceo Berrocal, V.; Muñoz Silva, M.; Nunez, I. N.; Matute, T. F.; Arce Medina, A.; Vidal, G.; Vidal Céspedes, C.; Dahlin, J. et al. Flapjack: Data Management and Analysis for Genetic Circuit Characterization. ACS Synthetic Biology 2021, 10, 183–191, Publisher: American Chemical Society.

(3) Galdzicki, M.; Clancy, K. P.; Oberortner, E.; Pocock, M.; Quinn, J. Y.; Rodriguez, C. A.; Roehner, N.; Wilson, M. L.; Adam, L.; Anderson, J. C. et al. The Synthetic Biology Open Language (SBOL) provides a community standard for communicating designs in synthetic biology. Nature biotechnology 2014, 32, 545–550.

(4) Roehner, N.; Beal, J.; Clancy, K.; Bartley, B.; Misirli, G.; Grunberg, R.; Oberortner, E.; Pocock, M.; Bissell, M.; Madsen, C. et al. Sharing structure and function in biological design with SBOL 2.0. ACS synthetic biology 2016, 5, 498–506.

(5) McLaughlin, J. A.; Beal, J.; Misirli, G.; Grünberg, R.; Bartley, B. A.; Scott-Brown, J.; Vaidyanathan, P.; Fontanarrosa, P.; Oberortner, E.; Wipat, A. et al. The synthetic biology open language (SBOL) version 3: simplified data exchange for bioengineering. Frontiers in Bioengineering and Biotechnology 2020, 8, 1009.

(6) Mante, J.; Abam, J.; Samineni, S. P.; Pötzsch, I. M.; Singh, P.; Beal, J.; Myers, C. J. Excel-SBOL Converter: Creating SBOL from Excel Templates and Vice Versa. bioRxiv 2022,

(7) Yanez Feliu, G.; Earle Gomez, B.; Codoceo Berrocal, V.; Muñoz Silva, M.; Nunez, I. N.; Matute, T. F.; Arce Medina, A.; Vidal, G.; Vidal Céśspedes, C.; Dahlin, J. et al. Flapjack: Data management and analysis for genetic circuit characterization. ACS Synthetic Biology 2020, 10, 183–191.

(8) McLaughlin, J. A.; Myers, C. J.; Zundel, Z.; Misirli, G.; Zhang, M.; Ofiteru, I. D.; Goni-Moreno, A.; Wipat, A. SynBioHub: a standards-enabled design repository for synthetic biology. ACS synthetic biology 2018, 7, 682–688.

(9) Galdzicki, M., et al. The Synthetic Biology Open Language (SBOL) provides a community standard for communicating designs in synthetic biology. Nature Biotechnology 2014, 32, 545–550.

(10) Mante, J.; Zundel, Z.; Myers, C. Extending SynBioHub’s Functionality with Plugins. ACS Synthetic Biology 2020, 9, 1216–1220.

(11) Vidal, G.; Vidal-Céspedes, C.; Rudge, T. J. LOICA: Integrating Models with Data for Genetic Network Design Automation. ACS Synthetic Biology 2022,

